# Sequencing, *de novo* assembly and annotation of the genome of the scleractinian coral, *Pocillopora acuta*

**DOI:** 10.1101/698688

**Authors:** Jeremie Vidal-Dupiol, Cristian Chaparro, Marine Pratlong, Pierre Pontarotti, Christoph Grunau, Guillaume Mitta

## Abstract

Coral reefs are the most divers marine ecosystem. However, under the pressure of global changes and anthropogenic disturbances corals and coral reefs are declining worldwide. In order to better predict and understand the future of these organisms all the tools of modern biology are needed today. However, many NGS based approaches are not feasible in corals because of the lack of reference genomes. Therefore we have sequenced, *de novo* assembled, and annotated, the draft genome of one of the most studied coral species, *Pocillopora acuta* (ex *damicornis*). The sequencing strategy was based on four libraries with complementary insert size and sequencing depth (180pb, 100x; 3Kb, 25x; 8kb, 12x and 20 kb, 12x). The *de novo* assembly was performed with Platanus (352 Mb; 25,553 scaffolds; N50 171,375 bp). 36,140 genes were annotated by RNA-seq data and 64,558 by AUGUSTUS (Hidden-Markov model). Gene functions were predicted through Blast and orthology based approaches. This new genomic resource will enable the development of a large array of genome wide studies but also shows that the *de novo* assembly of a coral genome is now technically feasible and economically realistic.

## Introduction

Coral reefs are the most divers marine ecosystem and the second one in terms of diversity after tropical rain forests (Knowlton et al. 2010). In addition to this ecological importance, it provides many ecosystem services, such as a direct access to food and economic-resources through fishing and tourisms (Done et al. 1996, Bryant et al. 1998). The physical and biological support of this ecosystem relies exclusively on one type of organisms, the hermatypic scleractinian coral. Worryingly, these sessile, colonial and symbiotic species are today threatened by an increasing number of various anthropogenic and natural disturbances (Hughes et al. 2003), inducing since the 80’s a worldwide decline of this ecosystem (Bellwood et al. 2004). As a consequence the research performed on these biological models has exponentially increased on all aspect of their biology.

Today, most methods studying variations at the transcriptomic, genetic or epigenetic level rely on next generation sequencing. The generalization of these approaches in model organisms such as in the mouse, drosophila or baker’s yeast has made it possible to achieve tremendous insights into the biology of these organisms leading to advances in numerous scientific domains (Koboldt et al. 2013). With the cost reduction of the NGS approach, this was also promised but is not yet fulfilled for non-model organisms because of the lack of reference genomes in many phyla or functionnal group. Among the anthozoa, reference genomes are available for the laboratory models *Nematostella vectensis* (Putnam et al. 2007) and *Aiptasia* sp. (Baumgarten et al. 2015). However, there is still few reference genomes for the ecologically important hermatypic corals. To date four species were sequenced and assembled: *Acropora digitifera* (Shinzato et al. 2011), *Pocillopora damicornis* (Cunning et al. 2018), *Stylophora pistillata* (Voolstra et al. 2017) and *Orbicela faveolata* (Prada et al. 2016). However, others are needed to widen the research effort on the future of biodiversity and corals adaptability to global changes.

Among the most studied coral is *Pocillopora acuta* (previously considered as a synonymous of *P. damicornis* until 2014 (Veron and Pichon 1976)). *P. acuta* is encountered on fringing reefs and sheltered areas in the entire Indopacific ocean (Veron 2000). *P. acuta* is also known to be very sensitive to many natural disturbances including coral bleaching (i.e. the symbiosis breakdown between the coral host and its micro-algae endosymbiont) (Loya et al. 2001) and diseases (Ben-Haim and Rosenberg 2002, Luna et al. 2007). It shows relatively good capacities of acclimatization to controlled condition and thus can be maintained in aquaria for decades where it present the advantage to be a fast growing species. These various features have led many scientists to use this coral as a model in a large variety of scientific fields such as population genetics (Adjeroud et al. 2013, Combosch and Vollmer 2015), integrative biology (Vidal-Dupiol et al. 2009, Vidal-Dupiol et al. 2011a), functional genomics (Traylor-Knowles et al. 2011), global changes impact (Vidal-Dupiol et al. 2013, Vidal-Dupiol et al. 2014), transgenerational acclimation (Putnam and Gates 2015), coral diseases (Ben-Haim Rozenblat and Rosenberg 2004, Vidal-Dupiol et al. 2011b), physiology (Richmond 1987, Stimson 1997) and host microbiota interactions (Bourne and Munn 2005) etc. which makes *P. acuta* a major scleractinian coral model.

Although transcriptomic resources are available (Traylor-Knowles et al. 2011, Vidal-Dupiol et al. 2011b) a reference genome for *P. acuta* is still lacking but could enable substantial advances in the understanding of various aspects of coral biology, ecology and evolution. In this context, the aim of the present study is to provide to the scientific community the draft genome sequence and annotation of *P. acuta*. To address this objective, the DNA from a single colony was sequenced using the Illumina technology and *de novo* assembled using Platanus assembler (Kajitani et al. 2014). The annotation of genes and repeats was performed. Gene prediction was conducted; firstly by an experimental annotation using previously published RNA-seq libraries and secondly, by an *ab initio* prediction based on a Hidden-Markov models. Finally, the putative function of each gene and its transcripts were identified by database searches and by an orthology based approach.

## Material and methods

### Biological material and DNA extraction

The *P. acuta* (Linnaeus, 1758) isolate used in this study was sampled in Lombok, Indonesia (Indonesian CITES Management Authority, CITES number 06832/VI/SATS/LN/2001-E; France Direction de l’Environnement, CITES number 06832/VI/SATS/LN/2001-I) and has been maintained in aquaria since the year 2001. Previously assigned to *Pocillopora damicornis* this isolate was reassigned to *P. acuta*. This assignation was based on the 840 based pair sequence of the ORF marker that enable to separate *P. acuta* and *P. damicornis* (Schmidt-Roach et al. 2014).

In order to avoid contaminations by the zooxanthellae for genome assembly *P. acuta* mini colonies (∼7 cm high, ∼6 cm diameter) used for DNA extraction were subjected to a menthol treatment inducing bleaching. Briefly, the colonies were placed in a four litter tank filled with seawater. Water motion was created using a submerged water pump (100L/h), temperature was maintained at 27°C and light adjusted to 75µmol/m^2^/s (PAR). The protocol for the menthol treatment was adapted from previous work (Wang et al. 2012). The first day, the corals were subjected to a concentration of menthol of 0.58mmol/L for 6h. After this exposure they were transferred to the coral nursery for a 18h recovery period. During the second day, the same protocol was applied (menthol treatment and recovery step). The third day, the coral were exposed again to the same treatment but only until the polyps were closed. Once achieved the corals were placed in the coral nursery for recovery while at the same time they lose zooxanthellae. This last step typically takes four to five days.

For DNA extraction, coral tissues of one bleached mini colony (∼7 cm high, ∼6 cm diameter) were harvested using an airpic in 50 mL of tissue extraction buffer (1M sucrose; 0.05 mM EDTA; 4°C). Then, the extract was centrifuged 10 min at 3000g (4°C) and the pellet was resuspended in the G2 buffer of the QIAGEN Genomic DNA kit. The rest of the protocol was performed according to the manufacturer’s instructions with the 100/G Genomic-tip. DNA quantity and quality were assessed by spectrophotometry (nanodrop), fluorescence (Qbit) and 0.5% agarose gel electrophoresis.

### Library preparation and sequencing

In order to facilitate the *de novo* assembly of the *P. acuta* genome, four different libraries were constructed and sequenced. Sequencing was performed on an Illumina Hiseq 2000 producing 100 bp paired-end reads. The first library was a shotgun (SG) library with an expected average insert size of 180 bp. It was sequenced at an estimated genome coverage of 100X. The second one was a long jumping distance (LJD) library with an average insert size of 3000 bp. It was sequenced for an estimated genome coverage of 25X. The two last one were LJD libraries with an expected average insert size of 8 and 20 kb, respectively. The sequencing coverage was estimated at 12X. Sequencing and library preparation were performed by the Eurofin company.

### *De novo* genome assembly

A stringent cleaning pipeline of the raw reads was applied to the four libraries. First, all reads of the four libraries were filtered in function of their quality. Only reads displaying a Phred quality score > 30 for 95% of its bases were kept for the analysis (FASTX-Toolkit). Secondly, the remaining reads were cleaned from any trace of adaptor using cutadapt program (Martin 2011). Thirdly, reads were interlaced/de-interlaced to separate singleton from paired reads and finally the reads from the SG library were subjected to a correction step using ErrorCorrection program from the SOAPdenovo2 package (Luo et al. 2012).

For the *de novo* assembly of *P. acuta* genome we used the multiple kmer assembler Platanus (Kajitani et al. 2014). For the contiguing step, only the paired read of the SG library were used. For scaffolding, the reads of all libraries were sequentially used from the shortest to the longest insert size (SG 180pb, LJD 3kb, LJD 8kb, LJD 20 kb). In a last step, gaps generated during scaffolding were closed using the paired reads of the SG libraries using the gap closer program included in the Platanus package. Once completed, the assembly quality was assessed by classical metrics (total assembly length, longest scaffold, N50 etc.) compiled in the program QUAST (Gurevich et al. 2013). In order to provide a functional validation of the assembly this quality assessment was also performed using the program CEGMA that looks for the presence of the 248 most conserved core eukaryotic proteins in the assembly (Parra et al. 2007). All scripts and parameters used in command line during the bioinformatics treatments are summarized in supplementary data 1, other tools were run on a local Galaxy instance.

### Structural annotation

As a first approach for genome annotation, exon/intron structures of genes were determined using RNA-seq data (experimental approach) previously published (Vidal-Dupiol et al. 2013, Vidal-Dupiol et al. 2014). The cleaned paired reads from our previous work were mapped against the scaffolds with a length above 5 kb using TopHat2 (Trapnell et al. 2009). All parameters were used with the default setting except for the mean and the standard deviation of the inner distance between pairs that were adapted to each specific library. Once mapped, the BAM file output obtained for each RNA-seq library was used to assemble the transcripts defined by TopHat2. This was done with Cufflinks with default parameters (Trapnell et al. 2010). Finally, all the transcript assembly generated (one per mapped RNA-seq library) were merged with Cuffmerge (Trapnell et al. 2010) using the default parameters.

Because our RNA-seq libraries potentially did not cover all putative transcripts in the genome, we performed as a second approach an *ab initio* gene prediction. This step was done using the AUGUSTUS web server (Stanke and Waack 2003) with the genome assembly filtered for scaffolds ≥ 5 kb and the Cuffmerge transcriptome as input file for the training and prediction steps. Genes were predicted on both strands with the option “predict any number of genes”.

### Functional annotation

In order to attribute a putative function to a maximum of the predicted transcripts but with the lowest probability of wrong annotation we applied two independent methods. The longest ORF of each putative transcripts was found using getorf (Rice et al. 2000), and the ones longer than 100 amino acids were selected for annotation. Then we used orthoMCL (Li et al. 2003) to identify potential orthologous sequences between our predicted ORF and; very well annotated genomes (*Homo sapiens* (Venter et al. 2001), *Mus musculus* (Chinwalla et al. 2002), *Caenorhabditis elegans* (Consortium 1998), *Danio rerio* (Howe et al. 2013), *Drosophila melanogaster* (Adams et al. 2000), *Saccharomyces cerevisiae* (Mewes et al. 1997) and *Strongylocentrotus purpuratus* (Sodergren et al. 2006)); or with a biological and an evolutionary interest with regard to the phylogenetic position of our organism (the cnidarians *Acropora digitifera* (Shinzato et al. 2011), *Hydra magnipapillata* (Chapman et al. 2010) and *Nematostella vectensis* (Putnam et al. 2007), the coral symbiont *Symbiodinium minutum* (Shoguchi et al. 2013) and the sponge *Amphimedon queenslandica* (Srivastava et al. 2010)). Orthology was considered significant when the *e*-value obtained was lower than 10^−5^. In this case and when available, the annotation of the ortholog(s) was transferred to the *P. acuta* sequence. In parallel to this approach we used Blast2GO (Conesa et al. 2005) version 2.4.2 to perform a semi-automated functional annotation of all putative transcripts using a set of similarity search tools (Conesa et al. 2005) : i) an initial annotation with BLASTX (against the non redundant NCBI database; *e*-value at 1 × 10^−3^); ii) a protein domain search using InterProscan; iii) an enzyme annotation using the *Kyoto Encyclopedia of Genes and Genomes* (Kanehisa and Goto 2000) (KEGG) enzyme database; and iv) assignment of a Gene Ontology term (Ashburner et al. 2000).

### Repeats annotation

Identification of the transposable elements (TEs) from the SINE family was performed as described previously (Baucom et al. 2009). Briefly, clustering was done by aligning candidates and manually extracting the families which were realigned and borders identified when possible. For the annotation of other TEs, the search using RepeatMasker (Tarailo-Graovac and Chen 2009) and the Repbase (Jurka et al. 2005) library on the assembled scaffolds did not yield any positive results. We therefore opted for another strategy which consists on recuperating all the reads that do not align to the assembled scaffolds using samtools (Li et al. 2009) and assemble them using RepArk (Koch et al. 2014) into contigs which will be searched for signatures of TEs. The RepeatMasker/Repbase combination did not give any results on the assembled contigs, therefore, we used RepeatModeler (Smit and Hubley 2010) to create a specific repeat library for the assembled contigs and used that library with RepeatMasker to search for TEs.

### Contamination level and genome heterozygosis

In order to evaluate the level of contamination by *Symbiodinium* sp. DNA that would be co-extracted with the coral DNA, we mapped the reads from the SG library on the genome of *S. kawagutii* (Lin et al. 2015) and *S. minutum* (Shoguchi et al. 2013). This was done with bowtie2 in single-end and sensitive end to end modes (Langmead and Salzberg 2012). In a second step, potential PCR duplicates generated during the amplification step of the gDNA-seq library preparation were removed using RmDup from the Sam tools package (Li et al. 2009) to decrease artefactual redundancy.

In order to evaluate the level of heterozygosis of the reference genome the raw reads of the SG library were mapped with bowtie2 on the *P. acuta* assembly. This was done in single end and sensitive end to end mode and as previously, PCR duplicates were removed with RmDup. Then, unique hits were filtered and used as input for variant calling using Varscan2 with default parameters (Koboldt et al. 2012).

## Results and Discussion

### Sequencing and de novo assembly

To enable the development of studies at the genome wide level and to strengthen the position of *Pocillopora acuta* as a coral model we sequenced and *de novo* assembled its genome. The sequencing strategy was based on 4 libraries with complementary insert sizes and sequencing depths (Table 1). The shotgun library (insert size 180 pb) yielded 237,589,976 paired reads of 100 bp each with an average Q score of 36.45 (93.71%>Q30). The corresponding 47,517 Mb produced correspond to an estimated genome coverage of 146X. The 3 kb LJD library has a measured average insert size of 2,479 bp. The sequencing yielded 109,242,128 paired read of 100 bp each with an average Q score of 32.31 (81.79%>Q30) this corresponds to 21,848 Mbp (genome coverage 67 x). The 8 kb LJD library has a measured average insert size of 7,825 bp. The sequencing yielded 44,399,583 paired read of 100 bp each with an average Qvalue of 31.98 (80.56%>Q30) which corresponds to 8,880 Mbp (genome coverage 27). The 20 kb LJD library has a measured average insert size of 32,325 bp. The sequencing yielded 99,813,316 paired read of 100 bp each with an average Qvalue of 30.76 (77.73%>Q30) which corresponds to 19,963 Mbp (genome coverage 61 x). In order to decrease the complexity of the dataset, the raw reads were quality filtered at Q30 for 90% of the read lenght, all remaining traces of adaptor were removed, and finally, sequencing error were corrected. Singleton were filtered and excluded from assembly.

**Table 1:** Data used for assembly

| Library | Average insert size<br>(measured; bp) | Paired-reads<br>(million) | Yield<br>(Mb) | Genome<br>coverage* | Average Q | Q>30( %) |
| --- | --- | --- | --- | --- | --- | --- |
| Shot-gun | 178 | 238 | 47,517 | 146 | 36.45 | 93.71 |
| LJD 3Kb | 2,479 | 109 | 21,848 | 67 | 32.31 | 81.79 |
| LJD 8Kb | 7,825 | 44 | 8,880 | 27 | 31.98 | 80.56 |
| LJD 20 Kb | 32, 325 | 100 | 19,963 | 61 | 30.78 | 77.73 |

\*For an estimated (kmer distribution) genome size of 325 Mb

The processed paired-reads were used as input for *de novo* assembly with Platanus assembler (contiging, scaffolding and gap closing). The assembly has resulted in 25,553 scaffolds greater than 1,000 bp and a N50 of 171,375 bp (the main assembly quality statistics are summarized in the Table 2). The longest scaffold reached 1,296,445 bp and the overall assembly is only 8% longer than the predicted genome size (352 Mb *vs* 325 Mb). These metrics are in the same order of magnitude than those classically obtained for other anthozoans (Putnam et al. 2007, Shinzato et al. 2011, Baumgarten et al. 2015). Considering the genome size, our prediction and assembly results stay in the anthozoa range delineated by *Aiptasia* sp. (the smallest; 260 Mb) and *Stylophora pistillata* (the largest; 434 Mb) genome (Baumgarten et al. 2015, Voolstra et al. 2017).

**Table 2:** Assembly statistics; all statistics are based on sequence of size ≥ 500 bp, unless otherwise noted

| Statistics | Contigs | Scaffolds |
| --- | --- | --- |
| Number (seq $\geq 0$ bp) | 196,891 | 168,465 |
| Number (seq $\geq 500$ bp) | 81,286 | 58,326 |
| Number (seq $\geq 1000$ bp) | 47,580 | 25,553 |
| Assembly length bp (seq $\geq 0$bp) | 336,691,489 | 352,019,984 |
| Assembly length bp (seq $\geq 500$bp) | 309,335,452 | 325,576,138 |
| Assembly length bp (seq $\geq 1000$bp) | 285,593,703 | 302,677,350 |
| Largest | 112,653 | 1,296,445 |
| N50 | 11,125 | 171,375 |
| NG50* | 10,244 | 171,375 |
| N75 | 3,847 | 16,625 |
| NG75* | 2,912 | 16,732 |
| N90 | 1,120 | 696 |
| NG90* | 818 | 521 |
| Number of N's per 100 kbp | 1.69 | 4697.98 |
| % Core eukaryotic genes (full length) | Not done | 84.3 |
| % Core eukaryotic genes (partial) | Not done | 93.15 |
| GC% | 37.84% |  |

\*based on the predicted genome size: 325 Mb

The GC% was equal to 37.84% and its distribution was unimodal. Since the GC % of anthozoa is around 37-39% (Shinzato et al. 2011) and around 44-46% in *Symbiodinium* sp. (Shoguchi et al. 2013, Lin et al. 2015) our results suggest a very low level of contamination by *Symbiodinium* DNA in our assembly (Sabourault et al. 2009). This very low level of contamination was confirmed through the mapping of the reads constituting the SG library on the *S. kawagutii* and *S. minutum* genome (Shoguchi et al. 2013, Lin et al. 2015). Indeed, only 0.02% and 0.03% of these reads were mapped on the genomes of *S. kawagutii* and *S. minutum* respectively. This results is strengthen by the very low number of contigs (one contig) included in an orthologs group containing Symbiodinium sequences only. These results confirmed that the menthol treatment was very efficient to induce a complete bleaching of *P. acuta* (Wang et al. 2012).

Finally, in order to evaluate the biological significance of this assembly we looked for the presence of 248 ultra-conserved core eukaryotic gene (COG) with the CEGMA package (Parra et al. 2007). In total, 84% of these COG were found in full length, and this number reach 93% when partial length similarities are included, showing the completeness of our assembly. These results are in the upper part of the range of values classically obtained from a *de novo* draft genome assembly performed with Illumina reads. For example, the draft genomes of the arthropods *Sarcoptes scabie* var. *hominis* and var. *suis* contained 98.79% of the 248 COG (Mofiz et al. 2016) while the draft genome of another arthropods, the chinese mitten crab *Eriocheir sinensis* contained 66.9% of these genes (Song et al. 2016).

### Structural and functional annotation

We had based this structural and functional annotation on experimental (RNA-seq) and *ab initio* approaches (Hidden-Markov model). Because these two approaches do not reach the same level of confidence we decided to keep the results of these two strategies separated. This will enable the users of these data to choose the experimental and/or *ab initio* gene predictions in function of the scientific question they will investigate.

As a first approach, genes were predicted using experimental data previously obtained. These RNA-seq libraries were chosen in order to cover a large range of physiological condition and include the response to the exposure to normal and high temperatures, virulent and non-virulent bacteria and acidification (Vidal-Dupiol et al. 2013, Vidal-Dupiol et al. 2014, Vidal-Dupiol et al. submitted). The paired reads of these libraries were used to predict gene and exon/intron structures on scaffolds with a size above or equal to 5 kb. This approaches has predicted 36,140 genes encoding 63,181 alternatively spliced transcripts (Table 3). These predicted transcripts were then used to train AUGUSTUS (Hidden-Markov model) for the *ab initio* gene prediction that had resulted in the identification of 64,558 predicted-genes encoding 79,506 alternatively spliced transcripts. The number of predicted genes using the experimental data is slightly higher to what was found for the two other sequenced symbiotic anthozoan (Shinzato et al. 2011, Baumgarten et al. 2015, Prada et al. 2016, Voolstra et al. 2017, Cunning et al. 2018). However the number of putative transcripts encoded by these genes are in agreement with what was classically obtained in Anthozoan transcriptome assembly (Meyer et al. 2009, Traylor-Knowles et al. 2011, Lehnert et al. 2012, Moya et al. 2012, Vidal-Dupiol et al. 2013, Shinzato et al. 2014, Kitchen et al. 2015). The differences between the number of gene predicted by the experimental and the *ab initio* approaches may result from difficulties encountered by AUGUSTUS rather than by TopHat2. Indeed, the mean exon length is 30% lower in the *ab initio* approach and this may result in an increasing gene number due to gene fragmentation. This hypothesis is confirmed by CEGMA that shows that the completeness of the transcriptome issued from the *ab initio* prediction is low with 56% of the 248 CEGs founds while 93% were found with the transcriptome generated by the experimental approach. In addition *ab initio* approaches are known to annotate some false genes such as pseudogenes and nonfunctional duplicated genes. However, both predictions can be usefull since genes specifically involved in some developmental process, larval stage, etc. may be represented in the *ab initio* annotation only, because genes pertaining to these physiological states were not found with the experimental annotation.

**Table 3:**
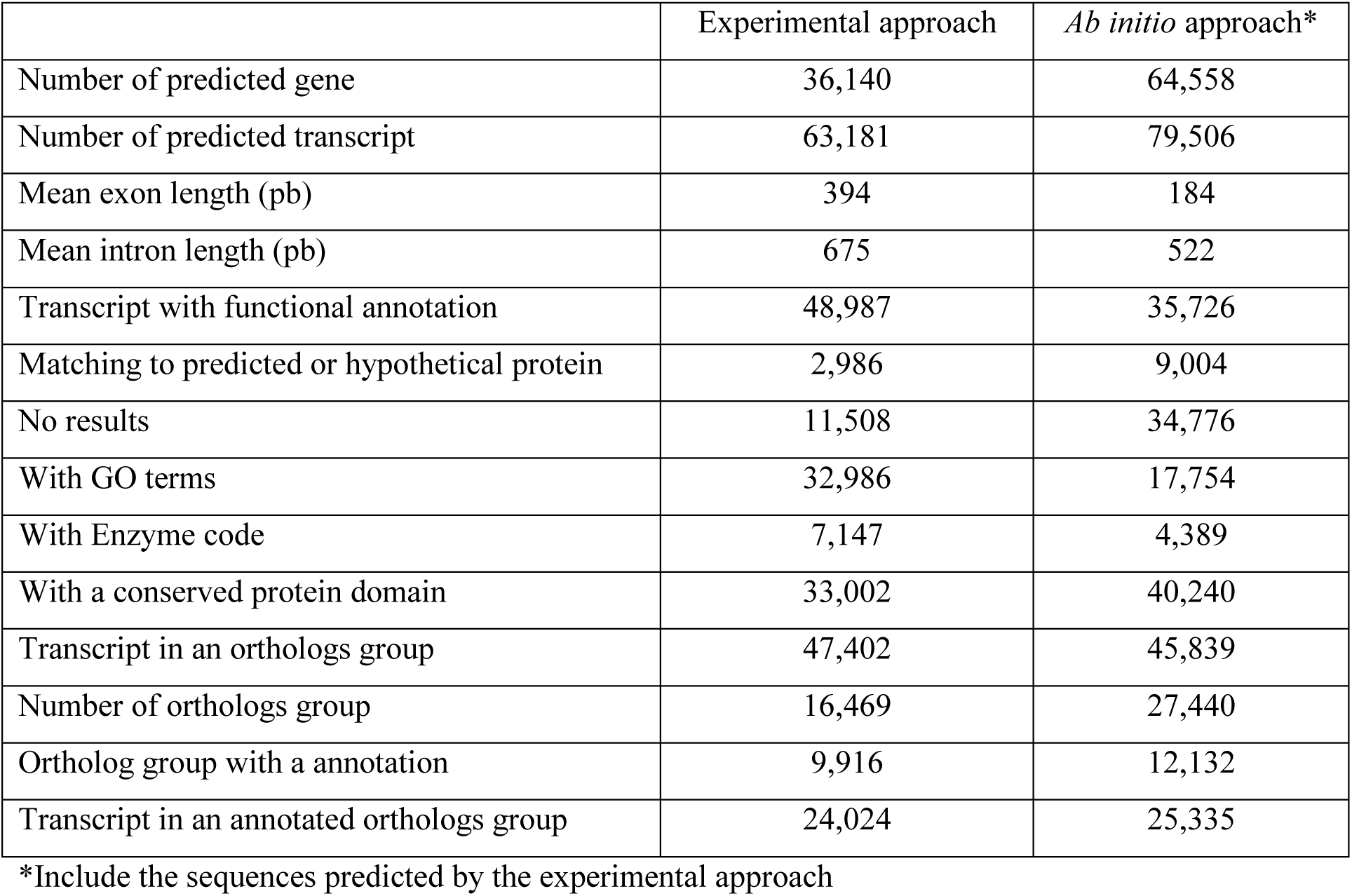
Structural and functional annotation summary results

The functional annotation was done using Blast2GO (Conesa et al. 2005) on the transcripts predicted from the RNA-seq and *ab initio* approaches, separately (Table 3). The BlastX performed on the non-redundant NCBI database returned the following results for the transcripts issued from the RNA-seq and *ab initio* approaches, respectively: i) 77.3% and 44.9% presented a significant similarity for a protein with a known function; ii) 4.6 % and 11.3% showed similarities for predicted proteins; iii) 18.1% and 43.8% did not returned any results; iv) GO-terms were then attributed to 52.2% and 22.3% of the transcripts; v) KEGG enzyme code was assigned to 11.3% and 5.5% of the sequences; and vi) conserved protein domains were detected in 52% and 50.6% of the transcript. Analysis of the RNA-seq based approached of GO-term repartition using WEGO (Ye et al. 2006) shows that our predicted transcripts enter in 6, 6 and 20 GO categories (GO level 2, for all level GO repartition, see the supplementary data 2) of the Cellular Component, Molecular Function and Biological Process root, respectively (Fig.1). For the RNA-seq based approach, orthoMCL had generated 16,469 groups of orthologs containing 47,388 transcripts (Table 3). This allowed to successfully annotating 24,024 transcripts with a high level of confidence (50.6% of those in clusters). For the *ab initio* approach, 27,440 groups of orthologs were created; they contain 45,839 transcripts and enable the annotation of 25,335 sequences. In comparison to what is usually obtained in transcriptome assembly of non-model invertebrates the number of transcripts showing significant similarities for protein with a putative function is high. Indeed, this rate fluctuates in general between 20 and 50% in many recent studies (Kitchen et al. 2015, Harney et al. 2016, McGrath et al. 2016). However in draft genome assembly this rates of annotation is higher and comparable to what we obtained (Shinzato et al. 2011, Baumgarten et al. 2015). This is probably due to a better sequencing coverage of the gene set in genome assembly rather than in transcriptome. Firstly, because genome *de novo* assembly need a higher sequencing coverage and secondly because in gDNA-seq the representation of each gene is theoretically equal while it is biased by the expression level in RNA-seq.

**Figure 1:**
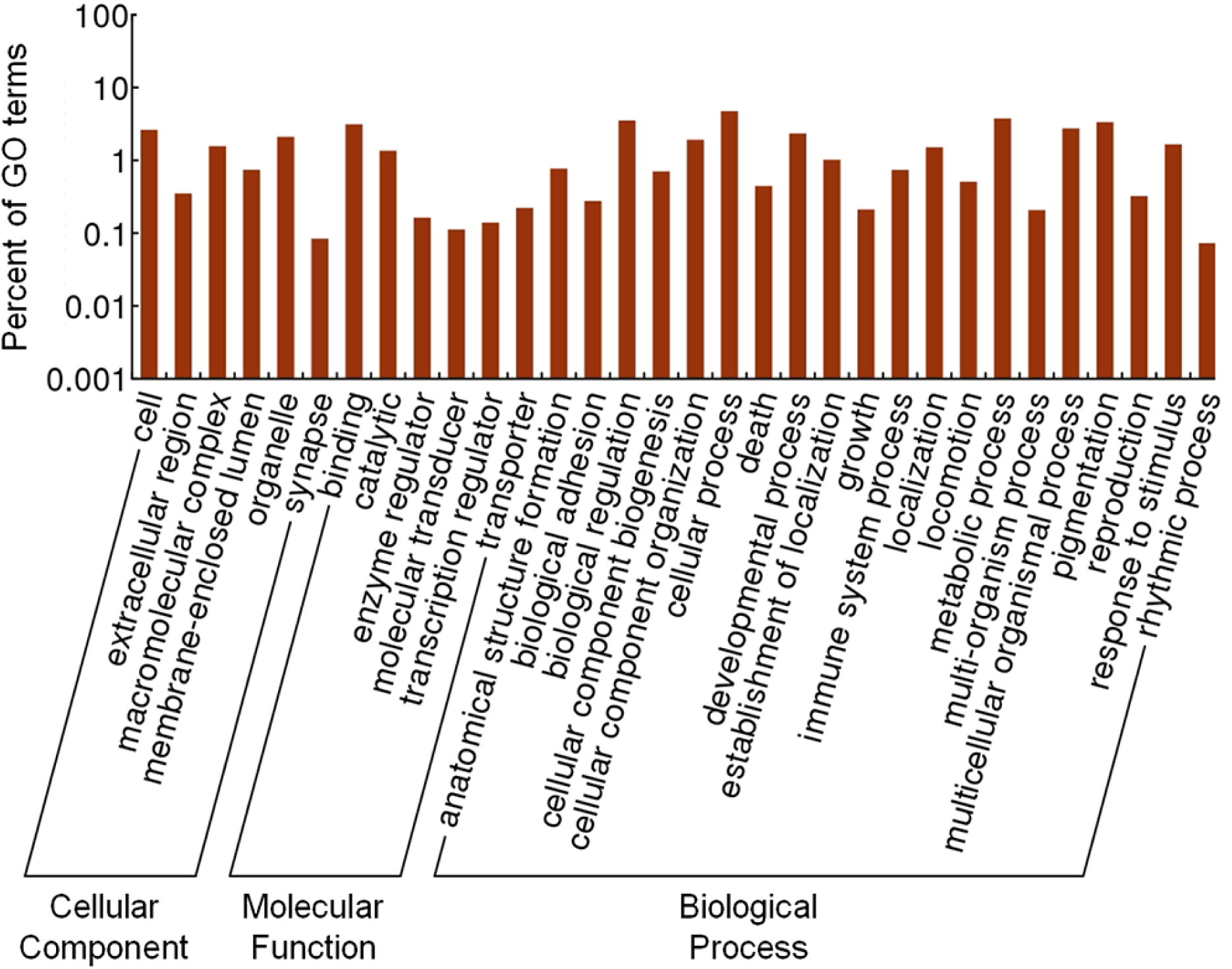
Distribution of GO terms belonging to the level 2 of the GO arborescence in the three main GO categories: Cellular Component, Molecular Function and Biological Process. Transcripts from the experimental annotation.

### Repeat content

The process of repeat annotation shows that at least 15.28% of *P. acuta* genome is composed of repeated sequences. This value is very close to what was found in the genome of *A. digitifera* with 13% of repeat content (Shinzato et al. 2011) and lower than in *Aiptasia* sp. with 26% (Baumgarten et al. 2015). In total, the four main superfamily of TEs (DNA transposons, SINE-, LINE- and LTR-retrotransposons) were found (Table 4) and they represent 18 families of transposons and 19 families of retrotransposons (supplementary data 3). Among them, seven are specific to *P. acuta* and 30 and 20 are shared with *Aiptasia* sp. And *A. digitifera*, respectively (Fig. 2). However, their occurrence is low in comparison to what was found in the two other symbiotic anthozoa (Table 4). Indeed, if 15.28% of the genome is composed of repeated sequences, the identified repeat sequences represent only 3.23% of this 15.36%. The remaining 11.98% correspond essentially to unidentified sequences (11.37%) in addition to satellites (0.03%), simple (0.9%), low complexity repeats (0.16%), and small RNA (0.06%). This high level of unidentified repeated sequences is not an exception. For example, in the *Aiptasia* genome, 63% of the repeat content corresponds to a unique unknown repeat (Baumgarten et al. 2015).

**Table 4:** *Pocillopora acuta*, repeat content

| Element (super family) | Number of occurrence | Number of bp covered | Per cent of genome coverage |
| --- | --- | --- | --- |
| DNA transposon | 27,081 | 3,708,267 | 1.05% |
| SINE retrotransposon | 8,504 | 11,051,79 | 0.31% |
| LINE retrotransposon | 45,381 | 4,352,179 | 1.24% |
| LTR retrotransposon | 3,457 | 838,736 | 0.24% |
| Unclassified | 364,626 | 40,009,286 | 11.37% |
| Small RNA | 2,751 | 198,347 | 0.06% |
| Satellites | 463 | 105,608 | 0.03% |
| Simple repeats | 62,511 | 3,182,003 | 0.90% |
| Low complexity | 11,794 | 576,390 | 0.16% |

**Figure 2:**
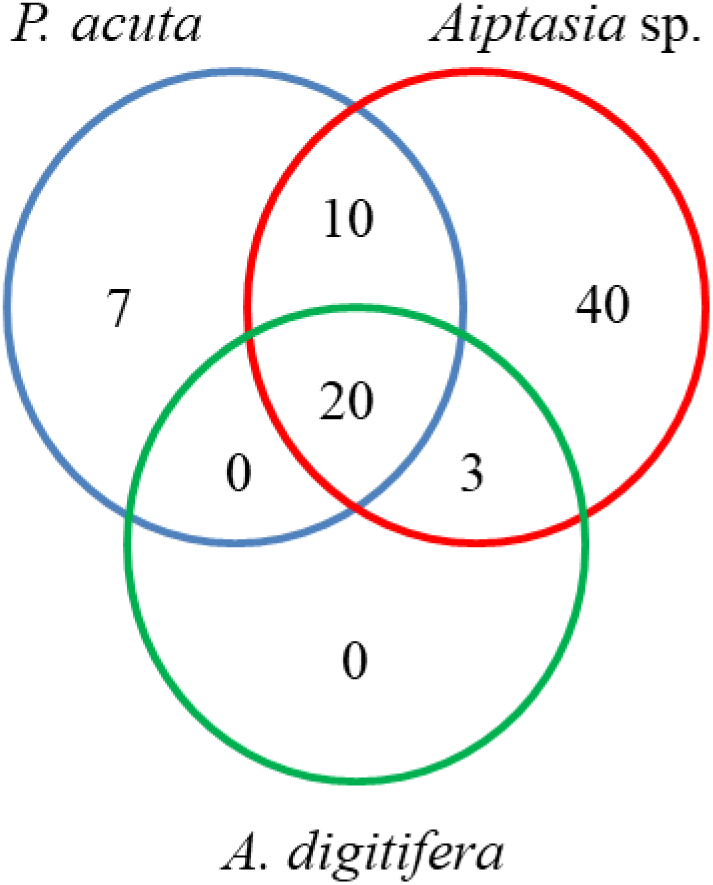
Venn diagram representing the number of transposable elements specific or shared between the three sequenced symbiotic anthozoa, the scleractinian corals *Acropora digitifera* and *Pocillopora acuta* and the anemonia *Aiptasia* sp. Transposable elements for *A. digitifera* and *Aiptasia* sp. were obtained from ^8,9^.

### Heterozygosis and SNP library

In order to evaluate the heterozygosis of the genome the reads from the SG library were re-mapped on the genome (Bowtie2) and variant were called with VarScan2. This analysis reveals the presence of 2,505,660 heterozygote single nucleotide polymorphisms (7.1 SNP/kb) and 321,295 indels. This level of heterozygosis in a single genome can be considered as very high even if we cannot exclude that a small proportion of these SNPs are false positive that can be due to sequencing or mapping error. However, if such a kind of data are absent from the anthozoan literature some comparisons are possible with other invertebrate phyla. In arthropods, SNP density can be as high as 16.5 SNP/kb in the butterflies from the genus *Lycaeides* (Gompert et al. 2010) and as low as 0.062 in the varroato-mite, *Varroa destructor* (Cornman et al. 2010). In these studies these results were obtained from dozen to hundreds of individuals, while in our study we have probably only one genotype per colony. This lead to believe that in the case of a populational study this density can significantly increase and reach very high values of SNP density. Alternatively, some of the polymorphism observed in our study can also be the results of intra colonial genetic variation, a phenomenon more and more observed and quantified in corals (Schweinsberg et al. 2014, Schweinsberg et al. 2015, Barfield et al. 2016) and that can be due to the accumulation of somatic mutations (Van Oppen et al. 2012) or to chimerism (Rinkevich et al. 2016).

### Cnidarian core proteome

The cnidarian core proteome was identified through the orthoMCL approach and its annotation. It is composed of 1,781 orthologous group of proteins, shared between the two scleractinian corals *P. acuta* and *A. digitifera*, the anemone *N*.*vectensis* (actinia) each of them belonging to the anthozoa class, and *H. magnipapillata* from the hydrozoa class (Fig.3). The GO term repartition (GO level 2) between this cnidarian core proteome and the entire proteome content of *P. acuta* is very close (Fig. 4) reflecting the absence of large functional gain or loss.

**Figure 3:**
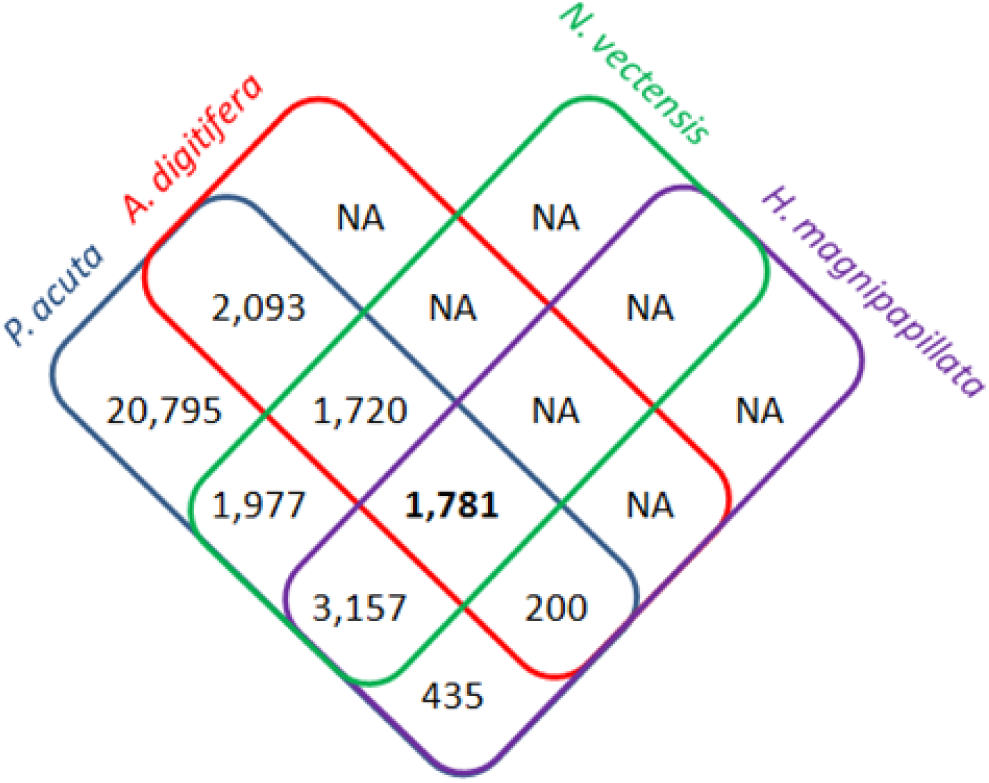
The cnidarian core proteome. This diagram represent the comparison between the protein content encoded in the genome of *P. acuta* and those of three other cnidarians: another scleractinia *A digitifera*; an actinia *N. vectensis*; and a hydrozoa *H. magnipapillata*. OrthoMCL was used to identify orthologous sequences between the predicted ORF (longer than 100AA) translated from the RNA-seq annotation and those annotated in these other genomes. The values in the diagram represent the number of group of orthologs created by orthoMCL (a group contains at least 2 sequences where at least one belongs to *P. acuta*). The value corresponding to the proteome restricted to *P. acuta* is composed by the number of ortholog groups containing *P. acuta* sequences only, plus the number of orphan sequences (sequences that are not included in a group). In total, this analysis include the 63,181 (47,402 are included in a group of ortholog, 15,779 are orphans) translated protein sequences annotated by RNA-seq in *P. acuta*, and 11,369, 10,636 and 7,278 sequences for *A digitifera, N. vectensis* and *H. magnipapillata*, respectively. NA signify not applicable: these values were not determined by this approach since the clustering was done to classify and annotate the proteome of *P. acuta*. Therefore, the proteins belonging to the other species and that do not has an ortholog in the genome of *P. acuta* could not be included in a ortholog group and so could not be taken into account by the analysis.

**Figure 4:**
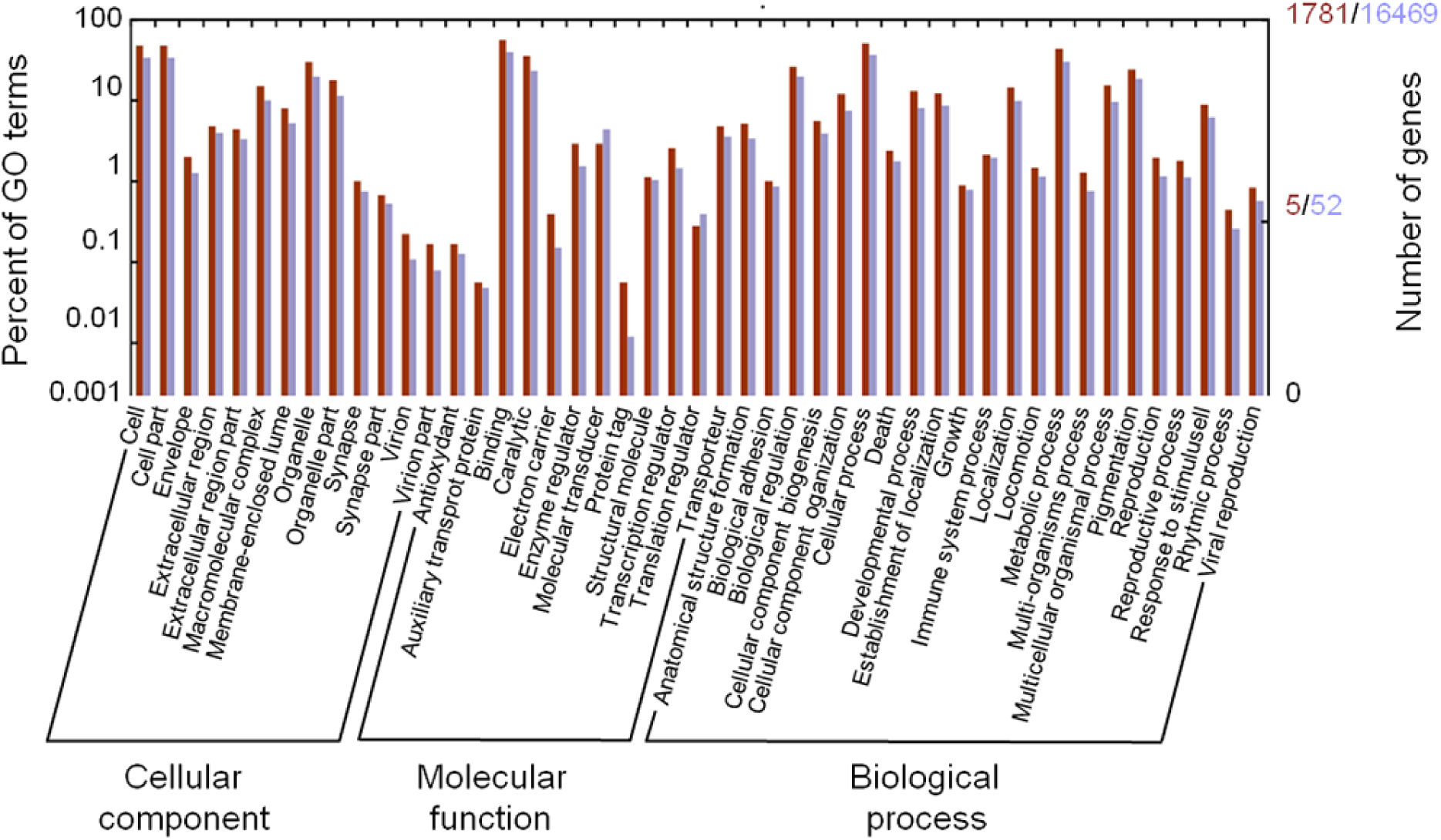
Distribution and comparison of GO terms included in the cnidarian core proteome (red) and the *P. acuta* proteome (blue). This GO terms belong to the level 2 of the GO arborescence and represent the three main GO categories: Cellular Component, Molecular Function and Biological Process. Transcripts from the experimental annotation.

### Conclusion

The draft genome assembly provided by this study constitutes a new coral reference genome and a new resource for the scientific community interested in cnidarians genomics approach (*sensus lato*). This will enable the development of a large array of genome wide studies that will lead to a better understanding of coral physiology, ecology and adaptability.

## Availability

The raw reads of each libraries were submitted to the NCBI Sequence Read Archive (accession numbers are: SRR4254617; SRR4254618; SRR4254619; SRR4254620) The draft genome and related annotation files can be downloaded using the following link http://ihpe.univ-perp.fr/acces-aux-donnees/

## Acknowledgements

This study was supported by the Agence Nationale de la Recherche through the Program BIOADAPT (ADACNI ANR-12-ADAP-0016-03) and the French-Israeli High Council for Science and Technology (P2R n u29702YG). The facilities of the Bio-Environnement platform (Perpignan, France) and the ABiMS platform (Roscoff, France) were used for the bioinformatics and molecular biology studies, the Aquarium facilities of the UMS 2348 were used for the sample preparation. The authors are indebted to Rayan Chikhi for his advices in bioinformatic during the assembly processes and to Sebastian Schmidt-Roach for his informations about the differences between *P. acuta* and *P. damicornis*.

